# Epigenetic dysregulation underpins tumorigenesis in a cutaneous tumor syndrome

**DOI:** 10.1101/687459

**Authors:** Helen R. Davies, Kirsty Hodgson, Edward Schwalbe, Jonathan Coxhead, Naomi Sinclair, Xueqing Zou, Simon Cockell, Akhtar Husain, Serena Nik-Zainal, Neil Rajan

**Author notes:** Corresponding authors: Neil Rajan, Institute of Genetic Medicine, Newcastle University, NE1 3BZ, UK. and Serena Nik-Zainal, Academic Department of Medical Genetics, University of Cambridge, Cambridge, UK. Email: Serena Nik-Zainal.

## Abstract

Patients with CYLD cutaneous syndrome (CCS; syn. Brooke-Spiegler syndrome) carry germline mutations in the tumor suppressor *CYLD* and develop multiple skin tumors with diverse histophenotypes ^1,2^. We comprehensively profiled the genomic landscape of 42 benign and malignant tumors across 13 individuals from four multigenerational families. Novel driver mutations were found in epigenetic modifiers *DNMT3A* and *BCOR* in 29% of benign tumors. Multi-level and microdissected sampling strikingly reveal that many clones with different *DNMT3A* mutations exist in these benign tumors, suggesting that intra-tumor heterogeneity is common. Integrated genomic and methylation profiling suggest that mutated *DNMT3A* drives tumorigenesis mechanistically through Wnt/ß-catenin pathway signaling. Phylogenetic and mutational signature analyses confirm the phenomenon of benign pulmonary metastases from primary skin lesions. In malignant tumors, additional epigenetic modifiers *MBD4*, *CREBBP, KDM6A* and *EP300* were mutated. We thus present epigenetic dysregulation as a driver in CCS tumorigenesis and propose this may account for the diverse histophenotypic patterns despite the paucity of mutations seen. These findings add novel dimensions to existing paradigms of cutaneous tumorigenesis and metastasis.

---

In human skin, benign tumors outnumber malignant tumors, yet genetic studies of these are limited ^3^. Rare inherited skin tumor syndromes such as CCS (Fig. 1a-b) offer an opportunity to address this knowledge gap and novel molecular insights into cancer can be gained. They may reveal unexpected driver mutations ^4^, highlight mechanisms that may be targetable with repurposed drugs developed for other cancers ^5^, or refine models of tumor growth and patterning. CCS patients develop multiple skin tumors named cylindroma, spiradenoma and trichoepithelioma, a histophenotypic spectrum of hair follicle-related tumors consistent with the hypothesis that they arise in hair follicle stem cells ^6,7^. These tumors occur both at sun-exposed and sun-protected sites. Infrequently, salivary gland tumors, pulmonary tumors ^8^, malignant transformation ^9^ and metastasis with lethal outcomes can occur.

**Fig. 1.**
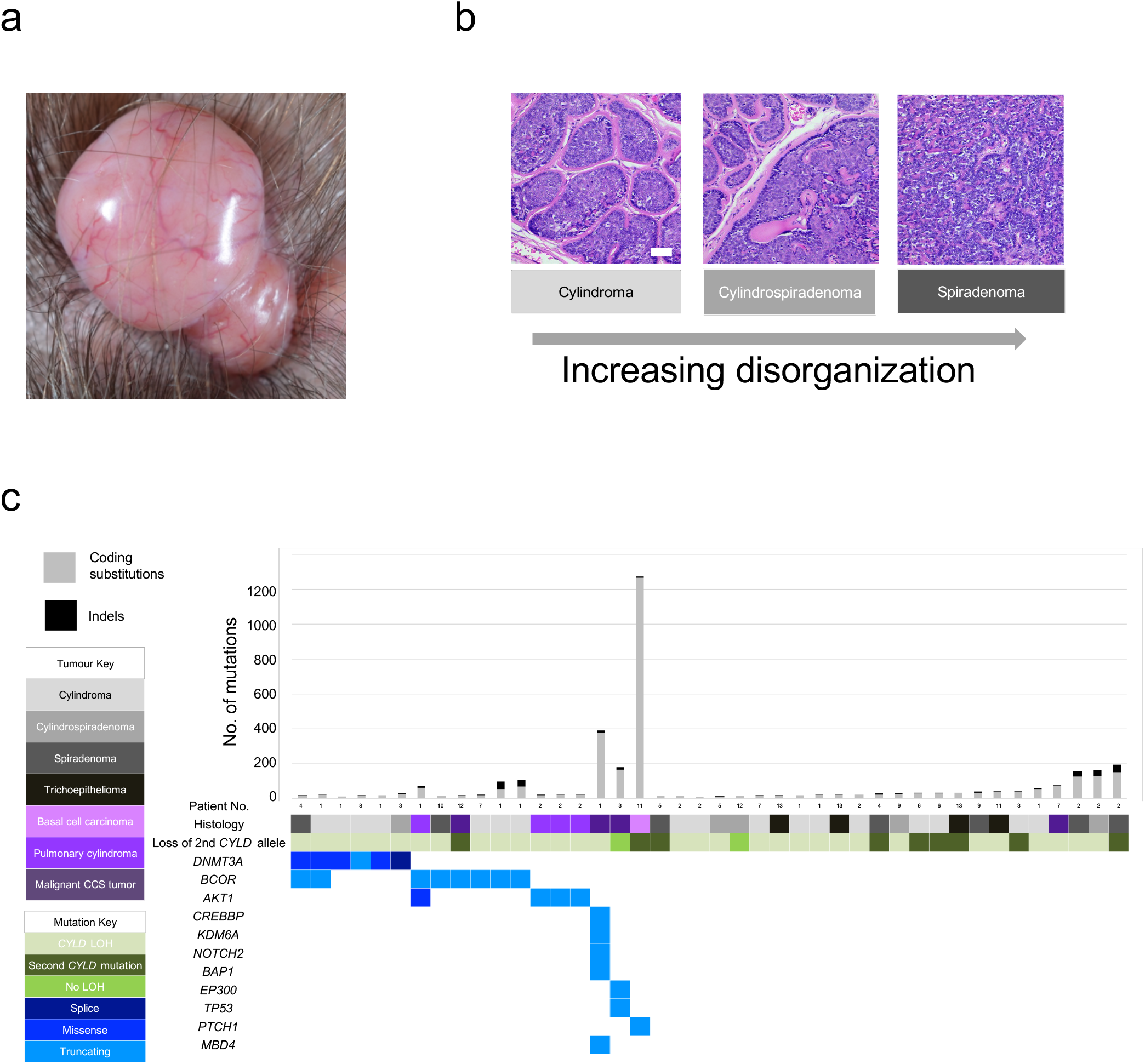
The mutational landscape of CYLD cutaneous syndrome. **a**, Benign but disfiguring cylindroma skin tumors of the scalp. **b**, Distinct histophenotypes of benign organised cylindroma and disorganised spiradenoma seen within the same sample, a frequent finding in CCS. (White scale bar=50um). **c**, Epigenetic modifiers are mutated in CCS tumors. Mutational burden is indicated in the bar graph with corresponding mutated genes shown below in the matrix. Matrix rows indicate mutated genes in each tumor and each matrix column represents a different sample (n=42).

*CYLD* encodes a ubiquitin hydrolase enzyme involved in deubiquitination of lysine 63 ^10,11^ and Met 1 linked ubiquitin chains ^12,13^. In CCS families, germline mutations occur within the catalytic domains of *CYLD* and are frequently truncating ^14^, predicting loss-of-function. Loss of the wild-type parental allele (loss of heterozygosity-LOH) of *CYLD* is demonstrated in the majority of inherited cylindromas, consistent with its role as a recessive cancer gene ^15^. Genetic analysis of sporadic cylindromas, rare in the general population, have highlighted the fusion transcript *MYB-NFIB* in a subset of tumors ^16^. Taken together with the recent finding of upregulated MYB in CCS tumors where *MYB-NFIB* fusions are absent, this supports MYB as a key downstream mediator of cylindroma pathogenesis following loss of *CYLD* ^17^. However beyond these drivers, CCS tumors studied using array-based comparative genomic hybridization (CGH) demonstrate a paucity of DNA aberrations, restricted to copy neutral LOH of *CYLD* ^15^, incongruent with the diverse histophenotypes seen within and across tumor samples.

Arguably, *CYLD* loss alone may be sufficient for tumorigenesis, *via* its role in negatively regulating oncogenic pathways; CYLD depletion using RNA interference first revealed its role in negatively regulating NF-κB signalling ^10,11,18^. Corroborating this, murine *CYLD* knockout models develop skin papillomas following chemical carcinogenesis that demonstrate increased expression of NF-κB target genes such as cyclin D1 mediated by dysregulation of BCL3 ^19^. Furthermore, CYLD has been shown to negatively regulate various oncogenic signalling pathways that are also relevant in hair development in embryogenesis, including Wnt ^20^, Notch and TGF-β ^6^.

In humans, recurrent loss of functional CYLD is reported in diverse cancers including: myeloma ^21^, leukaemia ^22,23^, hepatocellular carcinoma ^24^, neuroblastoma ^25^ and pancreatic cancer ^26^, consistent with its role as a tumor suppressor expressed ubiquitously in normal tissues. In CCS patients, increased Wnt signalling has been shown to be an oncogenic dependency in cylindroma and spiradenoma tumors ^7^. Histologically-organised cylindroma and histologically-disorganised spiradenoma represent extremes of a spectrum of histophenotype of the same tumor (Fig. 1b and **Supplementary Fig. 1a-c**). Transition from cylindroma to spiradenoma is associated with loss of expression of the negative Wnt signalling regulator Dickkopf 2 (*DKK2*)^7^. DNA methylation has been suggested as a mechanism to account for loss of expression of DKK2 ^7^, however comprehensive genomic and methylomic profiling of CCS tumors has not been performed. The inability of *CYLD* knockout mouse models to recapitulate the human phenotype of cylindroma tumors, has further limited characterisation of the genetic drivers in CCS ^27^.

## Results

### Biallelic loss of CYLD drives CCS tumors

Thus, to delineate the genomic landscape of CCS in humans, we studied DNA from 11 fresh frozen tumors using whole genome sequencing (WGS) in two directly-related patients who had been under clinical follow-up for 35 years (Pt. 1 and 2) (Fig. 1c and **Supplementary Table 1**). The average number of unique reads per tumor and normal sample for WGS was 374,496,607, generating 35.5 mean fold coverage for all samples. We detected on average 1,381 substitutions per tumor sample (average 0.44 mutations per MB), 72 small insertions and deletions (indels), and 1 rearrangement, using WGS. Biallelic mutations in *CYLD* were a recurrent driver mutation, and no *MYB*-*NFIB* fusions were found, consistent with previous studies (Fig. 1b)^15^. Tumors demonstrated neither recurrent structural rearrangements nor recurrent copy number aberrations (**Supplementary Fig. 2**).

To validate these findings, we studied a further 31 tumors from 12 patients of 3 additional genotyped pedigrees using whole exome sequencing, given the lack of large structural rearrangements. We confirmed that *CYLD* biallelic loss was independent for each sample, reinforcing that each tumor arose independently: Loss of the wild-type allele was observed either by LOH affecting 16q (31/42 tumors), or by a second mutation in *CYLD* (9/42), consistent with the loss of CYLD occurring across all benign and some malignant tumors in CCS.

### Selection of mutated epigenetic modifiers in benign CCS tumors

In addition to biallelic mutations in *CYLD*, we discovered multiple mutations in epigenetic modifiers DNA methyltransferase 3a (*DNMT3A)* (n=6) and BCL6 co-repressor (*BCOR*) (n=8) in 12 tumors (Fig. 2a, **Supplementary Fig. 3a and Supplementary Table 2**). In two tumors, both genes were mutated. *BCOR* mutations have also been reported to co-occur with *DNMT3A* in over 40% of BCOR-mutated cases of AML, suggesting a potential synergistic role in CCS tumorigenesis ^28^. Mutations in *DNMT3A* were predominantly missense mutations in the methyltransferase domain, but mutations in the zinc finger domains were also noted and have been reported previously in COSMIC (Fig. 2a) ^29^. Mutations in *BCOR* were predominantly frameshift mutations. Notably, different *DNMT3A* and *BCOR* driver mutations were seen in disparate tumors in patients 1 and 4, suggesting that convergent evolution drives tumorigenesis through epigenetic mechanisms in this cutaneous syndrome.

**Fig. 2.**
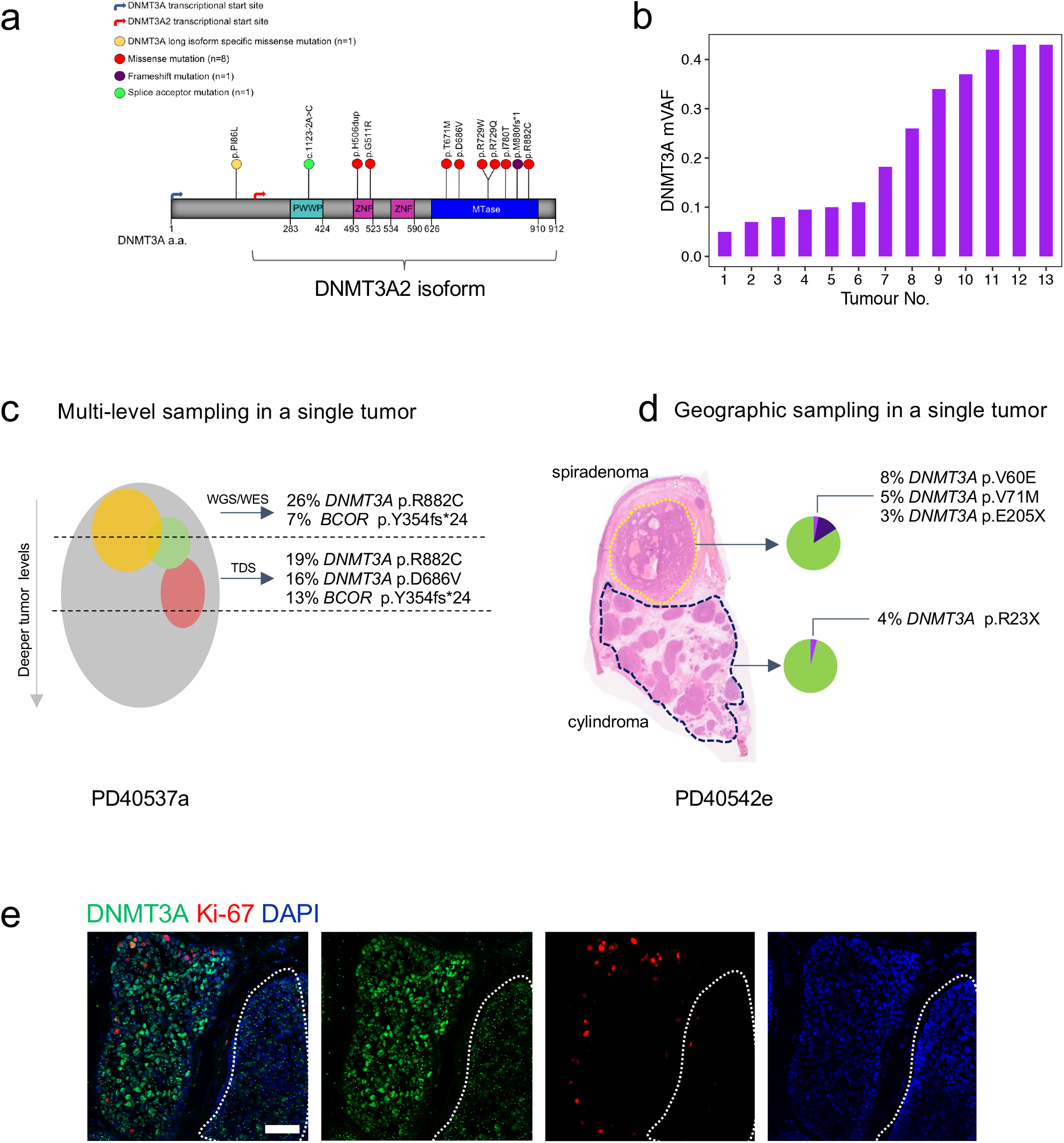
Intratumoral heterogeneity of *DNMT3A* mutation in CCS tumors. **a**, *DNMT3A* somatic mutation lollipop diagram for CCS tumors **b**, Spectrum of mutant variant allele fractions (VAF) of tumors in this study. **c**, Sampling of additional, deeper slices from a single tumor (PD40537a) reveals intratumoral heterogeneity of *DNMT3A* mutations (tumor indicated with grey sphere, intratumoral clones with coloured spheres). **d**, Geographic sampling of distinct histophenotypes (of cylindroma and spiradenoma) within a single tumor section (PD40542e) highlights marked clonal heterogeneity particularly of *DNMT3A* mutations. **e**, Protein expression of DNMT3A and Ki-67 is variable within a “cylinder” of CCS tumor and across cylinders (an adjacent outlined cylinder of cells showing of loss of DNMT3A expression).

Interestingly, variant allele frequencies of *DNMT3A* and *BCOR* mutations ranged from 0.05-0.42 – Fig. 2b), suggesting that intra-tumoral clonal heterogeneity may occur in these tumors. To explore this possibility, targeted deep sequencing (TDS; average coverage of >500x) of *DNMT3A* and *BCOR* was performed on additional material taken from further tissue sections of nine tumors studied above. This confirmed the presence of intratumoral heterogeneity of these putative driver mutations, with two distinct mutant clones or more found to co-occur within the same tumor in six samples (PD37330a,c,g,i, PD40536d, PD40537a) (Fig 2c and **Supplementary Table 1**).

To investigate whether *DNMT3A* mutational heterogeneity correlated with CCS tumor histophenotypes, we studied five tumors which contained intra-tumoral cylindroma and spiradenoma (Fig. 2d and **Supplementary Fig. 4a,b**). DNA was extracted from micro-dissected cylindroma and spiradenoma regions and TDS was performed. In three tumors, there was an identical *DNMT3A* mutation in both regions. In two tumors, there was heterogeneity between the histophenotypes, with private mutations in each regions, suggesting that multiple *DNMT3A* mutant clones of different sizes exist within tumors.

### Mutated DNMT3A2 epigenetically dysregulates Wnt/ß-catenin signalling

To explore the functional relevance of mutations in *DNMT3A* and *BCOR* in CCS tumors, RNA sequencing was performed in 16 tumors. This revealed increased expression of the short isoform of *DNMT3A*, called *DNMT3A2*, in 15 tumors compared to four perilesional skin controls (Fig. 3a). DNMT3A protein expression was also increased in CCS tumors compared to control skin and hair, and regions of heterogeneity were observed between islands of cylindroma (Fig. 2e and **Supplementary Fig. 5a,b**) in-keeping with reported mutational heterogeneity. *BCOR* was expressed at similar levels in both control and tumor tissue (**Supplementary Fig. 3b**).

**Fig. 3.**
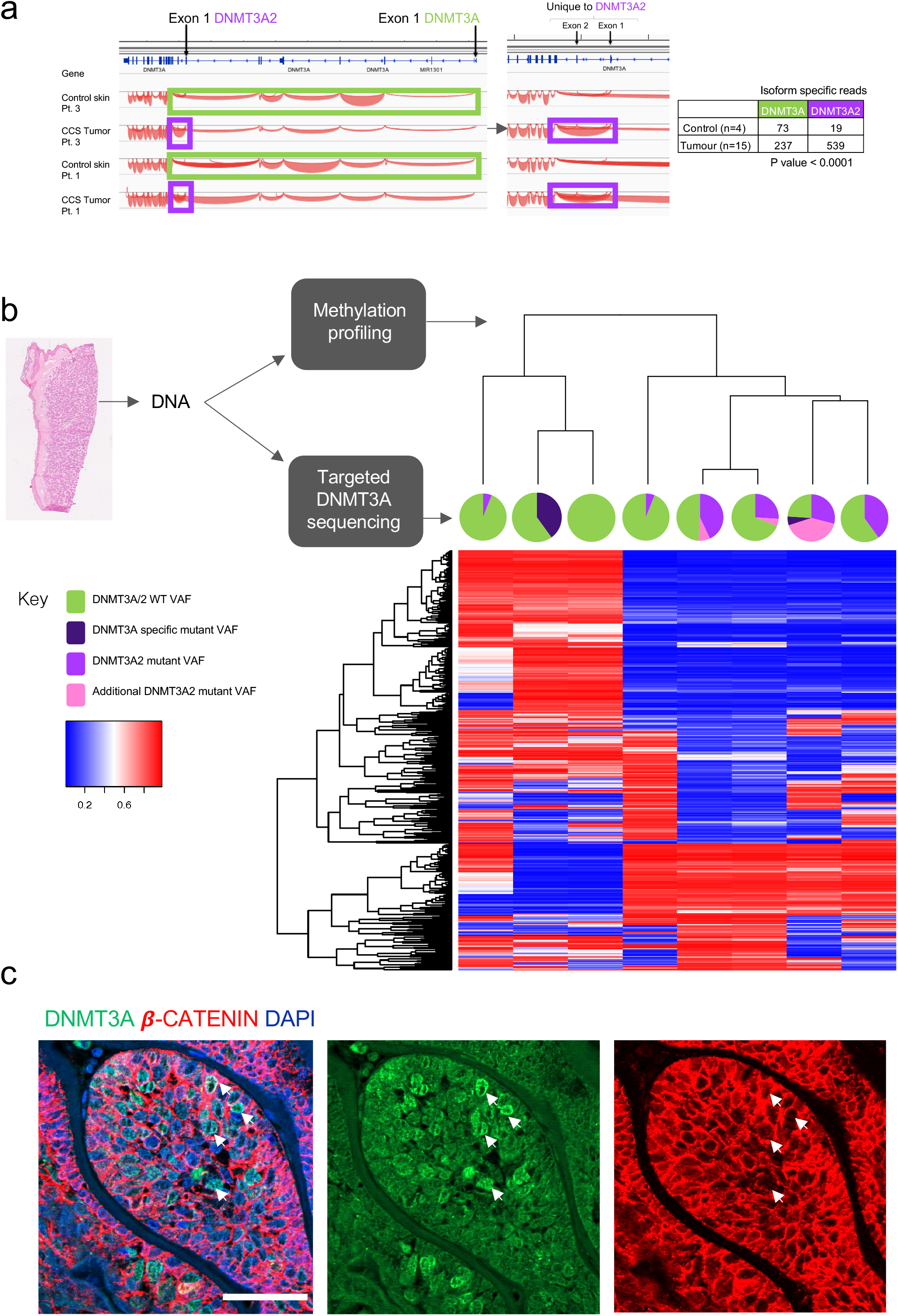
*DNMT*3*A*2 mutation alters methylation profiles of CCS tumors. **a**, RNA sequencing of 16 CCS tumors revealed that the short isoform of *DNMT3A*, *DNMT3A2*, is preferentially overexpressed in CCS. **b**, *DNMT3A2* mutant VAF is indicated as a pie chart format over a heatmap that displays most variable probes in 8 tumor samples that were subject to unsupervised clustering. **c**, DNMT3A and beta-catenin expression in CCS tumors demonstrate that DNMT3A expressing cells have low nuclear beta-catenin expression (white arrows).

To assess the impact of *DNMT3A* mutations on methylation patterns, eight samples genotyped by TDS were studied using genome-wide DNA methylation arrays. Unsupervised clustering of the 250 most variably methylated loci revealed a distinct cluster comprising five tumors with predicted *DNMT3A2* isoform specific mutations (“*DNMT3A2*-mutated”) (Fig. 3b). Comparison of these two clusters, revealed 1512 differentialy hypomethylated regions of contiguous probes in “*DNMT3A2*-mutated” tumors. Network analysis of these regions in “*DNMT3A2*-mutated” tumors identified the highest-ranked network to be functionally related to ß-catenin (p<1 × 10^−45^) (**Supplementary Fig. 6**). Taking this together with prior data that DNMT3A has been shown to preferentially methylate Wnt/ß-catenin signalling pathway genes ^30^, we assessed whether there was a relationship between expression of DNMT3A and expression of nuclear ß-catenin (Fig. 3c). Interestingly, this showed a co-relationship, where cells with reduced DNMT3A expression also demonstrated increased nuclear ß-catenin. These data support a mechanistic model where *DNMT3A2* isoform specific mutation selectively alters methylation on Wnt signalling genes, affecting transcriptional co-activator ß-catenin, which confers an oncogenic advantage in CCS tumor cells with known Wnt dependency.

### Malignant CCS tumors carry additional epigenetic modifier mutations

Malignant transformation although uncommon in CCS, is well-recognized. We studied five malignant CCS tumors: basal cell adenocarcinoma-low grade (BCAC-LG), malignant spiradenocarcinoma, atypical spiradenocarcinoma, poorly-differentiated adenocarcinoma and basal cell carcinoma (BCC) (**Supplementary Fig.7**) ^9^. The case of malignant spiradenocarcinoma (PD36119a) presented at the age of 80 in Patient 1. The tumor had a comparatively high number of coding substitutions (375 in the exome, corresponding to 8.4 per Mb), consisting largely of C>T transitions at CpG dinucleotides. This hypermutator phenotype has been reported previously in conjunction with germline methyl-binding domain 4 (*MBD4)* mutations^31^. Closer inspection confirmed a germline *MBD4* mutation in the patient, with concomitant loss of the wild-type parental allele in the tumor. Cascade screening revealed other family members that also carried this variant (**Supplementary Table 3**) although their tumors did not have biallelic *MBD4* loss and thus did not have the associated mutational signature. The observed burden and pattern of mutagenesis was consistent with *MBD4*’s role as a DNA glycosylase safeguarding the integrity of methylated CpGs from deamination. Notably, additional driver mutations detected included epigenetic modifiers, *KDM6A* and *CREBBP.* Tumor suppressors *NOTCH2* and *BAP1* were also noted to be driver mutations.

Poorly differentiated adenocarcinoma (PD40536c) has not been reported in CCS and presented on the breast of a female CCS patient at age 47. The patient had extensive staging scans, mammograms and biopsies of breast cylindromas, and has been followed up for 3 years with no evidence of a non-cutaneous primary tumor. This tumor had driver mutations in *TP53* and the epigenetic modifier *EP300*. Strikingly, this did not demonstrate LOH for *CYLD*. The BCAC-LG (PD40545a) tumor demonstrated a frameshift mutation in *BCOR*. The atypical spiradenocarcinoma (PD40540a) did not show any changes apart from *CYLD* LOH. The BCC (PD45044c) demonstrated a *PTCH* driver mutation and *CYLD* LOH, consistent with genetic features of BCC ^32^. It also demonstrated the highest number of coding substitutions (1287) in our cohort, comprising the UV signature, in contrast to benign trichoepithelioma also arising on the face of the same patient.

### Mutation analysis tracks origin of pulmonary cylindromas to the skin

To investigate mutational mechanisms that may give rise to the mutations detected in CCS patients, we compared the mutational signatures in tumors with identical histological types at intermittently sun-exposed and typically sun-protected sites ^33^ (Fig. 4a). Two tumors from the torso demonstrated substitution signature 7 (n=2; PD37331a,i) consistent with UV exposure. By contrast, we did not find evidence of signature 7 and found the presence of mutational signatures 1 (associated with deamination of methylated cytosines) and 5 (unknown etiology) in sun-protected tumors from pubic and perianal sites (n=4; PD37330c,e,g and PD37331c) and some intermittently sun-exposed tumors from the breast and torso (n=2). We surmise that in CCS, additional mechanisms other than UV are relevant to development of skin cancer.

**Fig. 4.**
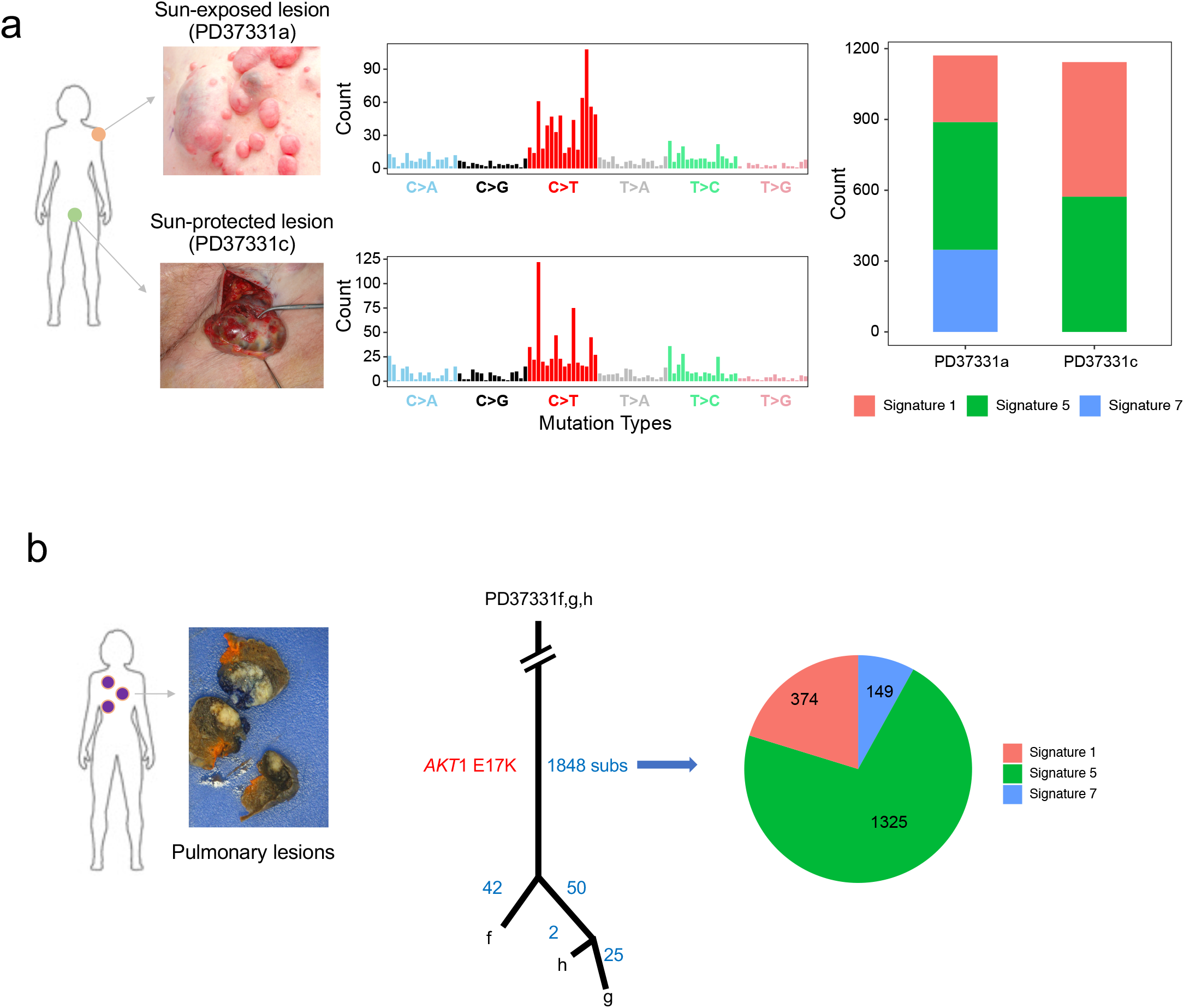
UV signature analysis reveals distinct mutational mechanisms in skin and tracks origin of lung tumors. **a**, Examples of intermittently sun-exposed and sun-protected CCS tumors demonstrate differing mutational profiles. Mutational signature analysis reveals UV-related signature 7 in sun-exposed tumors only. **b**, In one patient with three pulmonary lesions, a phylogenetic analysis reveals 1,848 mutations were shared in common and showed a UV signature. Hence these benign pulmonary lesions had a common origin, likely-sun-exposed skin.

We next investigated the concept of “benign metastases” seen in some patients with CCS, who develop multiple pulmonary cylindromas without typical features of malignancy ^8^. We studied four pulmonary cylindromas that had benign histological features from patients 1 and 2, who were both ex-smokers. They did not have evidence of lymph node disease, hepatic or bone metastases (Fig. 4b). Tumor phylogenetic analysis revealed that multiple pulmonary lesions from patient 2 shared 1,848 substitutions suggesting that these geographically-separated lesions that seeded in the lung, had a common origin (Fig. 4c). We found that the UV mutation signature 7 was present in the shared mutations, and thus tracked the origin of these pulmonary lesions to sun-exposed skin. Lastly, we found recurrent, E17K *AKT1* oncogenic mutations in multiple lung cylindromas in each patient, and in both patients independently as well. This is interesting for two reasons: First, although the numbers are small, this suggests that *AKT1* mutations likely arose prior to seeding in the lung. The *AKT1* mutations may confer lung tissue tropism for cylindromas. Second, this recurrent *E17K AKT1* mutation is clinically-relevant and targetable. As drugs have been developed to target *AKT*1 mutations in a diverse range of solid tumors ^34^, this finding further creates therapeutic opportunities for this limiting secondary complication of CCS.

## Discussion

This work delineates the mutational landscape of CCS. It highlights the presence of distinct *DNMT3A* and *BCOR* mutations in different tumor sites of the same patient (inter-tumor heterogeneity) and different geographic sites within the same tumor (intra-tumor heterogeneity), which suggests strong convergent evolution (**Supplementary Fig. 8**) towards epigenetic dysregulation in this orphan disease where no medical treatments are available. Due to the clinical interest in mutated epigenetic modifiers in leukemia, strategies used to target *DNMT3A* mutant hematological malignancies may be relevant to CCS ^35^. The accessibility of CCS skin tumors lend themselves to direct drug delivery, which may be an attractive route avoiding systemic side effects, as suggested by the methodology of a recent early phase clinical trial in CCS ^5^.

In addition, we reveal mutational synergy between *CYLD* loss and epigenetic modifiers such as *DNMT3A*, through NF-κB and Wnt signalling pathways. This may have broader relevance for common BCC, where *CYLD* is transcriptionally repressed ^36^, and Wnt/ ß-catenin signalling is pathogenic ^37^. Notably, a recent study reported the presence of mutations in *DNMT3A* (10%) and *BCOR* (18%) in the exomes of 293 BCCs; functional studies are required to investigate the clinical significance of this finding ^32^. Finally, we uncategorically demonstrate that the multiple benign pulmonary lesions in this syndrome have a clonal, cutaneous ancestral origin – reinforcing the concept of benign metastases as a clinical phenotype.

## Materials and Methods

### Patients and samples

Retrospective review of the case notes and radiological data of 15 genotyped *CYLD* mutation carriers that were under follow up between 1 July 2013 and 1 July 2017 was performed. Skin and lung samples were obtained from patients with signed consent, and details of samples are shown in Table 1. Research ethics committee approval was obtained for this work (REC Ref: 06/Q1001/59; 08/H0906/95+5).

#### Histology and immunohistochemistry

Histological assessment was performed following standard H+E staining and in conjunction with a dermatopathologist (A.H.). Immunofluorescent labelling with antibodies against DNMT3A, Beta Catenin and Ki 67 was carried out as described previously (1). Briefly, tissue sections from snap frozen skin tumor biopsies were fixed, blocked and then probed overnight at 4°C with primary antibodies. Antibodies against DNMT3A (#3598) and Ki-67 (#9449) were obtained from Cell Signalling, USA. Beta Catenin antibody (#610153) was obtained from BD Transduction USA. Secondary fluorescent antibodies (Alexa Fluor #111-545-144 488-conjugated Goat-Anti Rabbit and #115-585-146 594-conjugated Goat-Anti Mouse) were applied the following day and visualized with a fluorescent microscope (Zeiss Axioimager Z2, with Apotome 2 – Carl Zeiss, UK).

#### Accession codes

Bam files for whole genome sequencing were deposited in European Genome-phenome Archive under the following accession number: EGAD00001004573. FASTQ files for whole exome sequencing were deposited under accession numbers: ***EGA-BOX 1146***. FASTQ files for RNA sequencing were deposited under accession number: ***EGA-BOX 1146***. FASTQ files for targeted deep sequencing were deposited under accession number: ***EGA-BOX 1146***. Methylation array files were deposited under accession number: ***EGA-BOX 1146*** DNA was extracted from 12 cases along with corresponding normal tissue and subjected to paired-end WGS on an Illumina HiSeq X Ten, as described previously^***33,38***^. DNA for WES was extracted from blood and cyrosections of snap frozen tissue, and in 5 cases, from FFPE tissue (PD37330h, PD40536c, PD40540a, PD40545a and PD40545c). 42 WES library samples were prepared using the Illumina Nextera DNA exome kit, prior to being sequenced on a S2 flowcell on an Illumina Novaseq machine. Three WES samples were enriched using the SureSelect Human All ExonV6+UTR and 100 base paired-end sequencing performed on an Illumina Hiseq 2500 genome analyzers. For WES sequence depth was on average 255***-**fold*. Resulting BAM files were aligned to the reference human genome (GRCh37) using Burrows-Wheeler Aligner, BWA BWA-0.7.16a (r1181). Mutation calling was performed as described previously ^***33***^. Briefly, CaVEMan (Cancer Variants through Expectation Maximization:http://cancerit.github.io/CaVEMan/) was used for calling somatic substitutions. Indels in the *tumor* and normal genomes were called using a modified Pindel version 2.0 (http://cancerit.github.io/cgpPindel/) on the NCBI37 genome build. Structural variants were discovered using a bespoke algorithm, BRASS (BReakpoint AnalySiS; https://github.com/cancerit/BRASS) through discordantly mapping paired-end reads followed by *de novo* local assembly using Velvet to determine exact coordinates and features of breakpoint junction sequence. All mutations were annotated according to ENSEMBL version 75.

### ASCAT copy number analysis

Allele-specific copy number analysis of tumors analyzed by WGS was performed using ASCAT (v2.1.1) as described previously ^33^. ASCAT takes non-neoplastic cellular infiltration and overall tumor ploidy into consideration, to generate integer-based allele-specific copy number profiles for the tumor cells. Copy number values and estimates of aberrant tumor cell fraction provided by ASCAT were in put into the CaVEMan substitution algorithm for WGS. In addition, ASCAT segmentation profiles were used to establish the presence of loss of heterozygosity across *CYLD* and relevant mutated cancer driver genes.

### Identification of driver mutations

Somatic mutations present in known cancer genes (Cancer gene census https://cancer.sanger.ac.uk/census) were reviewed to identify those which were likely to be driver mutations. Mutations were deemed to be potential driver mutations if they were consistent with the type of mutations found in a particular cancer gene; that is, inactivating mutations in tumor suppressor genes (including nonsense, frameshift, essential splice site mutations and recurrent missense) and recurrent mutations in dominant oncogenes. Recurrent mutations were determined by reference to reported mutation frequency in the COSMIC database (https://cancer.sanger.ac.uk/cosmic).

### Mutational signature analysis

The contributions of substitution signatures for WGS samples were determined as follows: The substitution profile is described as a 96-channel vector. For each mutation, of which there are 6 substitution classes of C>A, C>G, C>T, T>A, T>C and T>G, the flanking 5’ and 3’ sequence context is taken into account giving a total of 96 channels. A given set of mutational signatures was fitted into the mutational profile of each sample to estimate the exposure of each of the given signatures in that sample. The fitting algorithm (Degasperi *et al*., in preparation) detects the presence of mutational signatures with confidence, using a bootstrap approach to calculate the empirical probability of an exposure to be larger or equal to a given threshold (i.e. 5% of mutations of a sample). Here, we first used 30 COSMIC signatures (https://cancer.sanger.ac.uk/cosmic/signatures) to fit into each sample, and then chose the first three signatures with highest confidence, which are signature 1, 5 and 7, to do the final fitting.

For highly mutated malignant samples (the spiradenocarcinoma (PD36119a) and the BCC (PD40544c)), the mutation burden was orders of magnitude higher than other non-malignant tumors that were exome-sequenced. We were able to use cosine similarity between the overall 96 channel profile and COSMIC signature to confirm the presence of particular mutational signatures in the relevant sample. The cosine similarity between each malignant sample and the suspected COSMIC signature was high: For PD36119a, cosine similarity to COSMIC signature 1 was 0.92 and for PD40544c cosine similarity to the UV light signature, COSMIC signature 7, was 0.98.

### Targeted sequencing

The Truseq Myeloid panel (Illumina) was used to sequence *DNMT3A* and *BCOR* in 18 samples accordance with the manufacturers protocol. A 20 pM library of the PhiX genome was added to achieve a 5% PhiX spike-in. This library was loaded onto a Miseq flow cell (600 cycles V3) for sequencing (Illumina, San Diego, CA, USA). Data was analysed using BWA (v.0.7.15) to align reads to the reference sequence and Samtools used as a variant caller. Variant calls that passed strict filtering thresholds (“Filter”=PASS and “Qual”=100) were included for the deep sequencing on sections in additional levels and in new samples. For five samples (PD37330k, PD37331k, PD37331m, PD40542e and PD 40536e – Supplementary Figure S4) where intratumoral clonal variation was studied across distinct histophenotypic regions, variant call thresholds were relaxed, and all non-synonymous variants called were confirmed by visualizing aligned read data using Integrated Genomics Viewer (IGV; v2.3). These variants were included if aligned reads supported the variant calls.

### Transcriptomic analyses

RNA was extracted from 16 tumor samples and 4 control samples and stranded preparation was performed using the Illumina stranded mRNA kit as previously described^5^. Libraries were prepared and sequenced using an Illumina Hiseq 2500, giving 15 million paired end reads per sample which were 100 bp in length. FASTQ files were aligned using the splice aware aligner program STAR to generate alignment files ^39^. The read counts for each sample file were counted using the R package Subread ^40^. Differential gene expression analysis was carried out using the package DeSeq2 ^41^.

### Methylation assay

We assessed genome-wide DNA methylation in seven tumor samples with the Illumina Methylation EPIC microarray (Illumina, San Diego, CA, USA). DNA methylation assays were performed as per the standard manufacturer’s protocol by MWG (Aros – Denmark). Methylation array processing, functional normalization^42^ and quality control checks were implemented using the R package minfi ^43^. Differentially methylated probes were identified using minfi. Differentially methylated regions spanning multiple probes were identified using bumphunter ^44^; these regions were visualized using Gviz ^45^.

## Supporting information

Supplementary Table 4. A full list of all mutations detected.

## Acknowledgments

We are indebted to the patients and families who took part in this study. We are grateful for helpful discussions with Debbie Hicks, Nick Reynolds, Joris Veltman and Muzlifah Haniffa.

## Author contributions

N.R, E.S, H.D, K.H, N.S, X.Z, S.C, S.N-Z, A.H. contributed to the experiments, scientific hypotheses, data analysis and compiling of the manuscript. J.C and A.H contributed to the experiments and data analysis. N.R and S.N-Z designed the experiments N.R, H.D, S.N-Z, wrote the manuscript.;

## Data and materials availability

Sequencing data from this study are stored in the EGA and are accessible at accession numbers EGAD00001004573 and ega-box-1146 (submission in process)

## Supplementary Information

Supplementary Figures 1-8

Supplementary Tables 1-4

## References

1. Rajan, N. et al. Tumor mapping in 2 large multigenerational families with CYLD mutations: implications for disease management and tumor induction. Arch Dermatol 145, 1277–84 (2009).

2. Bignell, G.R. et al. Identification of the familial cylindromatosis tumour-suppressor gene. Nat Genet 25, 160–5 (2000).

3. Marino-Enriquez, A. & Fletcher, C.D. Shouldn’t we care about the biology of benign tumours. Nat Rev Cancer 14, 701–2 (2014).

4. Tomlinson, I.P. et al. Germline mutations in FH predispose to dominantly inherited uterine fibroids, skin leiomyomata and papillary renal cell cancer. Nat Genet 30, 406–10 (2002).

5. Danilenko, M. et al. Targeting Tropomyosin Receptor Kinase in Cutaneous CYLD Defective Tumors With Pegcantratinib: The TRAC Randomized Clinical Trial. JAMA Dermatol 154, 913–921 (2018).

6. Rajan, N. & Ashworth, A. Inherited cylindromas: lessons from a rare tumour. Lancet Oncol 16, e460–e469 (2015).

7. Rajan, N. et al. Transition from cylindroma to spiradenoma in CYLD-defective tumours is associated with reduced DKK2 expression. J Pathol 224, 309–21 (2011).

8. Brown, S.M. et al. Inherited pulmonary cylindromas: extending the phenotype of CYLD mutation carriers. Br J Dermatol 179, 662–668 (2018).

9. Kazakov, D.V. et al. Morphologic diversity of malignant neoplasms arising in preexisting spiradenoma, cylindroma, and spiradenocylindroma based on the study of 24 cases, sporadic or occurring in the setting of Brooke-Spiegler syndrome. The American journal of surgical pathology 33, 705–719 (2009).

10. Brummelkamp, T.R., Nijman, S.M., Dirac, A.M. & Bernards, R. Loss of the cylindromatosis tumour suppressor inhibits apoptosis by activating NF-kappaB. Nature 424, 797–801 (2003).

11. Trompouki, E. et al. CYLD is a deubiquitinating enzyme that negatively regulates NF-kappaB activation by TNFR family members. Nature 424, 793–6 (2003).

12. Draber, P. et al. LUBAC-Recruited CYLD and A20 Regulate Gene Activation and Cell Death by Exerting Opposing Effects on Linear Ubiquitin in Signaling Complexes. Cell Rep 13, 2258–72 (2015).

13. Kupka, S. et al. SPATA2-Mediated Binding of CYLD to HOIP Enables CYLD Recruitment to Signaling Complexes. Cell Rep 16, 2271–80 (2016).

14. Nagy, N., Farkas, K., Kemeny, L. & Szell, M. Phenotype-genotype correlations for clinical variants caused by CYLD mutations. Eur J Med Genet 58, 271–8 (2015).

15. Rajan, N. et al. Dysregulated TRK signalling is a therapeutic target in CYLD defective tumours. Oncogene 30, 4243–60 (2011).

16. Fehr, A. et al. The MYB-NFIB gene fusion-a novel genetic link between adenoid cystic carcinoma and dermal cylindroma. J Pathol 224, 322–7 (2011).

17. Rajan, N. et al. Overexpression of MYB drives proliferation of CYLD-defective cylindroma cells. J Pathol 239, 197–205 (2016).

18. Kovalenko, A. et al. The tumour suppressor CYLD negatively regulates NF-kappaB signalling by deubiquitination. Nature 424, 801–5 (2003).

19. Massoumi, R., Chmielarska, K., Hennecke, K., Pfeifer, A. & Fassler, R. Cyld inhibits tumor cell proliferation by blocking Bcl-3-dependent NF-kappaB signaling. Cell 125, 665–77 (2006).

20. Tauriello, D.V. et al. Loss of the tumor suppressor CYLD enhances Wnt/beta-catenin signaling through K63-linked ubiquitination of Dvl. Mol Cell 37, 607–19 (2010).

21. Jenner, M.W. et al. Gene mapping and expression analysis of 16q loss of heterozygosity identifies WWOX and CYLD as being important in determining clinical outcome in multiple myeloma. Blood 110, 3291–300 (2007).

22. Braun, T. et al. Targeting NF-kappaB in hematologic malignancies. Cell Death Differ 13, 748–58 (2006).

23. Hahn, M. et al. Aberrant splicing of the tumor suppressor CYLD promotes the development of chronic lymphocytic leukemia via sustained NF-kappaB signaling. Leukemia 32, 72–82 (2018).

24. He, G. & Karin, M. NF-kappaB and STAT3 - key players in liver inflammation and cancer. Cell Res 21, 159–68 (2011).

25. Kobayashi, T., Masoumi, K.C. & Massoumi, R. Deubiquitinating activity of CYLD is impaired by SUMOylation in neuroblastoma cells. Oncogene 34, 2251–60 (2015).

26. Taniguchi, K. & Karin, M. NF-kappaB, inflammation, immunity and cancer: coming of age. Nat Rev Immunol 18, 309–324 (2018).

27. Rajan, N. & Ashworth, A. Inherited cylindromas: lessons from a rare tumour. The lancet oncology 16, e460–9 (2015).

28. Tiacci, E. et al. The corepressors BCOR and BCORL1: two novel players in acute myeloid leukemia. Haematologica 97, 3–5 (2012).

29. Forbes, S.A. et al. COSMIC: exploring the world’s knowledge of somatic mutations in human cancer. Nucleic Acids Res 43, D805–11 (2015).

30. Rinaldi, L. et al. Loss of Dnmt3a and Dnmt3b does not affect epidermal homeostasis but promotes squamous transformation through PPAR-gamma. Elife 6, 1350 (2017).

31. Rodrigues, M. et al. Outlier response to anti-PD1 in uveal melanoma reveals germline MBD4 mutations in hypermutated tumors. Nature Communications, 1–6 (2018).

32. Bonilla, X. et al. Genomic analysis identifies new drivers and progression pathways in skin basal cell carcinoma. Nat Genet 48, 398–406 (2016).

33. Nik-Zainal, S. et al. Landscape of somatic mutations in 560 breast cancer whole-genome sequences. Nature 534, 47–54 (2016).

34. Hyman, D.M. et al. AKT Inhibition in Solid Tumors With AKT1 Mutations. J Clin Oncol 35, 2251–2259 (2017).

35. Rau, R.E. et al. DOT1L as a therapeutic target for the treatment of DNMT3A-mutant acute myeloid leukemia. Blood 128, 971–81 (2016).

36. Kuphal, S. et al. GLI1-dependent transcriptional repression of CYLD in basal cell carcinoma. Oncogene 30, 4523–30 (2011).

37. Yang, S.H. et al. Pathological responses to oncogenic Hedgehog signaling in skin are dependent on canonical Wnt/beta3-catenin signaling. Nat Genet 40, 1130–5 (2008).

38. Alexandrov, L.B., Nik-Zainal, S., Wedge, D.C., Campbell, P.J. & Stratton, M.R. Deciphering signatures of mutational processes operative in human cancer. Cell Rep 3, 246–59 (2013).

39. Dobin, A. et al. STAR: ultrafast universal RNA-seq aligner. Bioinformatics 29, 15–21 (2013).

40. Liao, Y., Smyth, G.K. & Shi, W. The Subread aligner: fast, accurate and scalable read mapping by seed-and-vote. Nucleic Acids Res 41, e108 (2013).

41. Love, M.I., Huber, W. & Anders, S. Moderated estimation of fold change and dispersion for RNA-seq data with DESeq2. Genome Biol 15, 550 (2014).

42. Fortin, J.P. et al. Functional normalization of 450k methylation array data improves replication in large cancer studies. Genome Biol 15, 503 (2014).

43. Aryee, M.J. et al. Minfi: a flexible and comprehensive Bioconductor package for the analysis of Infinium DNA methylation microarrays. Bioinformatics 30, 1363–9 (2014).

44. Jaffe, A.E. et al. Bump hunting to identify differentially methylated regions in epigenetic epidemiology studies. Int J Epidemiol 41, 200–9 (2012).

45. Hahne, F. & Ivanek, R. Visualizing Genomic Data Using Gviz and Bioconductor. in Statistical Genomics: Methods and Protocols (eds. Mathé, E. & Davis, S.) 335–351 (Springer New York, New York, NY, 2016).

