## Supplementary Information for "Epigenetic dysregulation underpins tumorigenesis in a cutaneous tumor syndrome"

**This PDF file includes:**

Supplementary Figures 1-8
Supplementary Tables 1-4

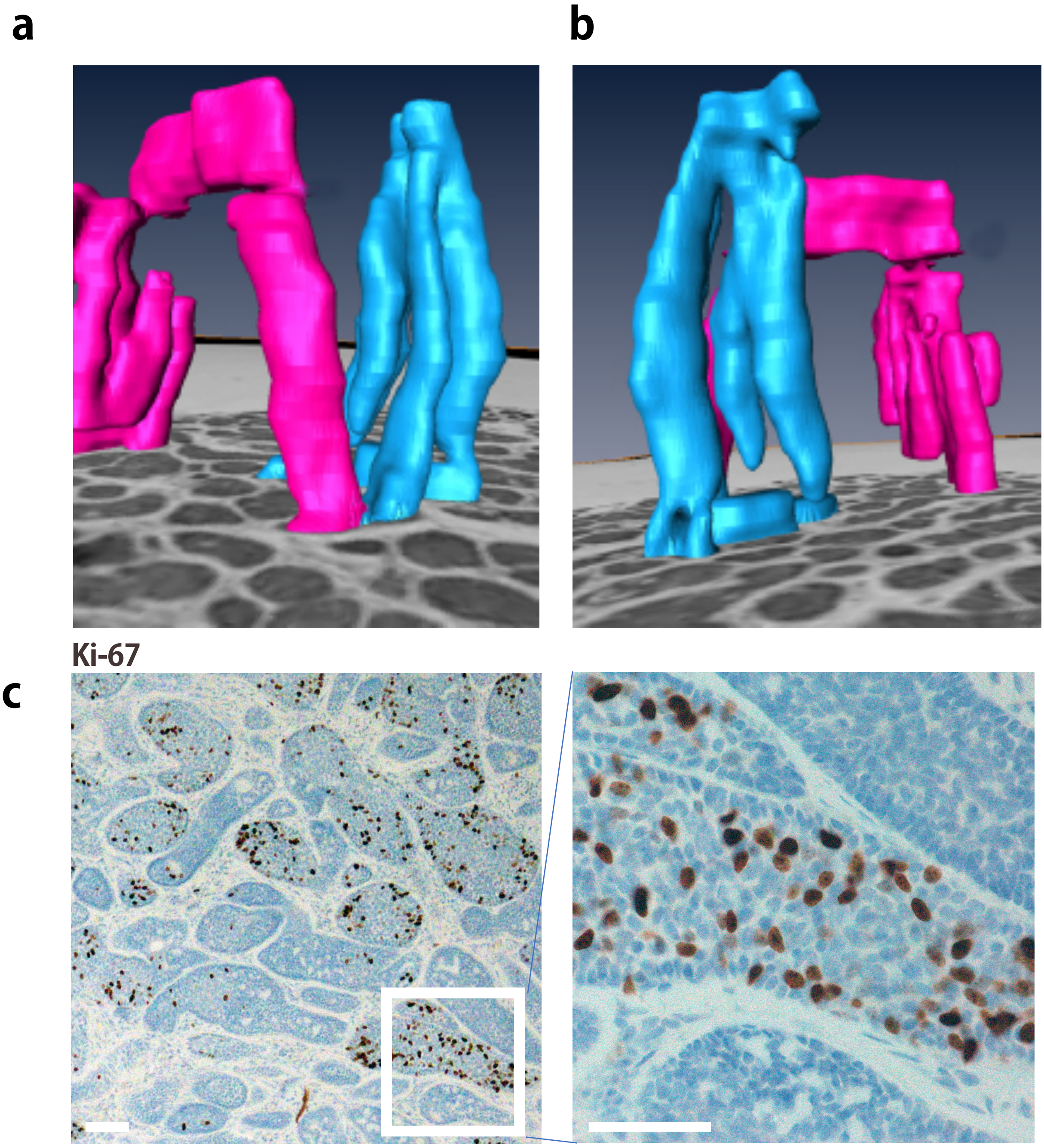


**Supplementary Figure 1. Histological patterns of growth of CCS tumors. a,b** Three-dimensional reconstruction of tumors from serial sections *in silico* highlight adjacent islands of tumor as distinct cylindrical compartments indicated in blue and red, with (**c)** a range of states of proliferation as indicated by expression of the proliferation marker Ki-67.

**a**

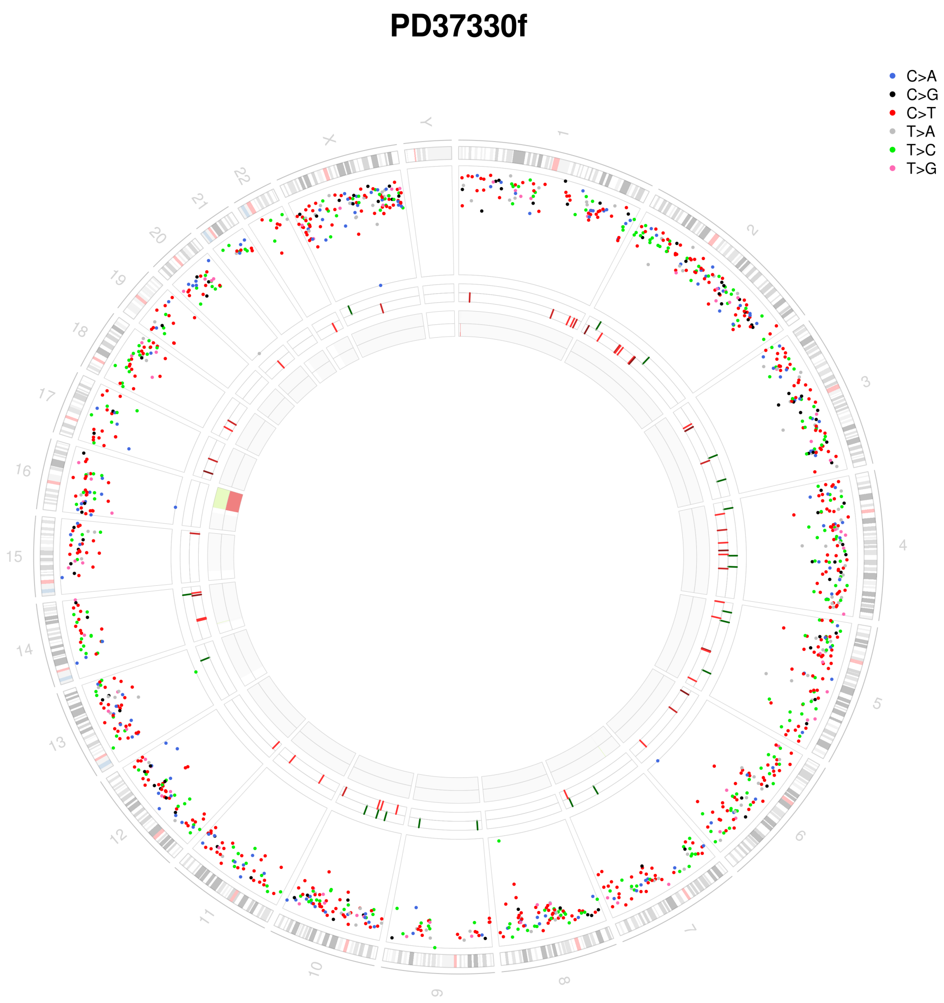


**Supplementary Figure 2. Loss of heterozygosity affecting 16q is a recurrent feature in CCS tumors. a** Whole genome circos plot depicting from outermost rings heading inwards: Karyotypic ideogram outermost. Base substitutions next, plotted as rainfall plots ($log^{10}$ intermutation distance on radial axis, dot colours: blue, C>A; black, C>G; red, C>T; grey, T>A; green, T>C; pink, T>G). Ring with short green lines, insertions; ring with short red lines, deletions. Major copy number allele ring (green, gain), minor copy number allele ring (red, loss).

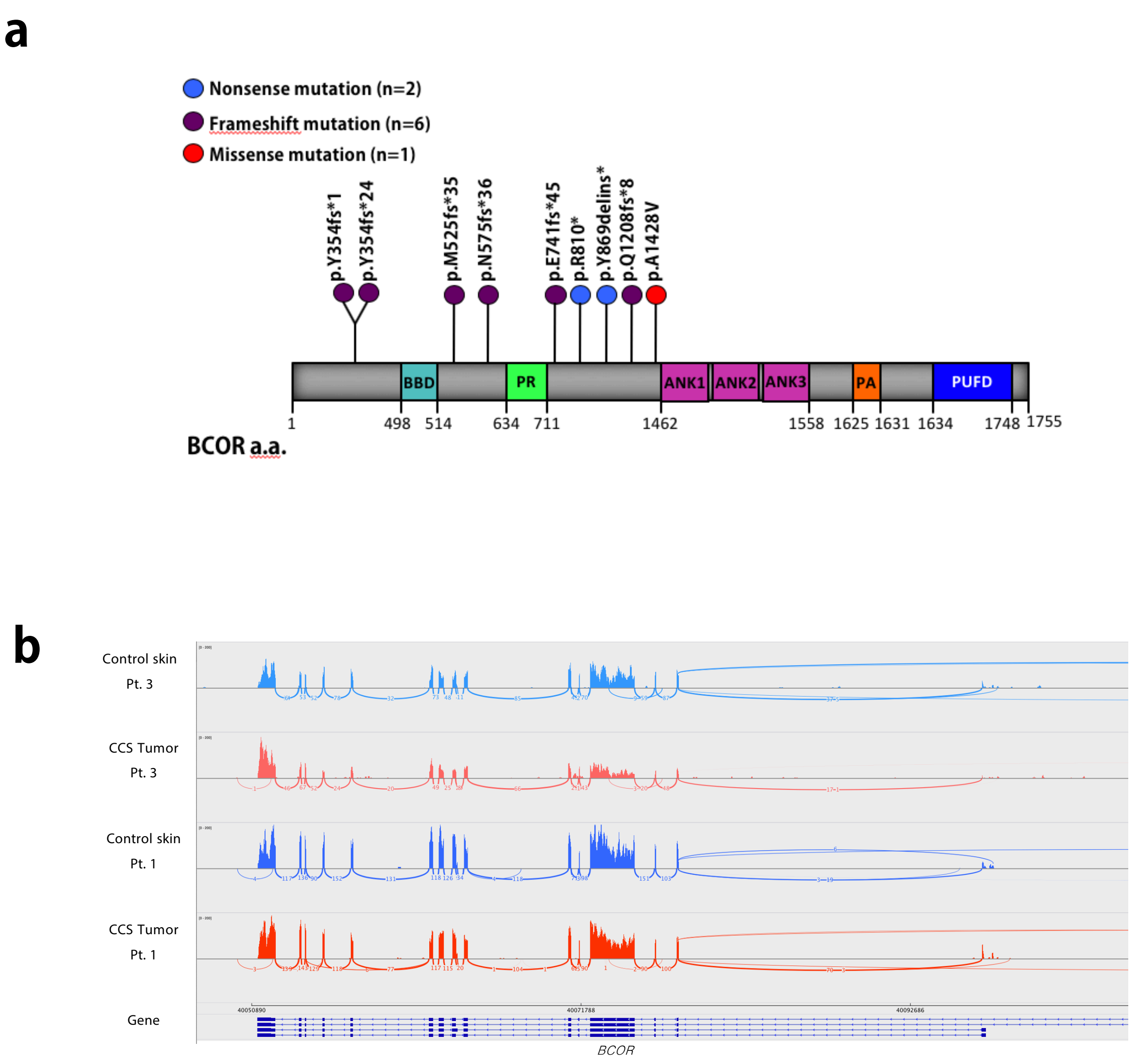


**Supplementary Figure 3. Mutation spectrum of *BCOR* and expression in CCS.** **a** Lollipop diagram indicating the distribution of mutations in *BCOR*. **b** RNA sequencing data demonstrating indicative expression of *BCOR* in 2 CCS tumors and matched patient controls.

**a**

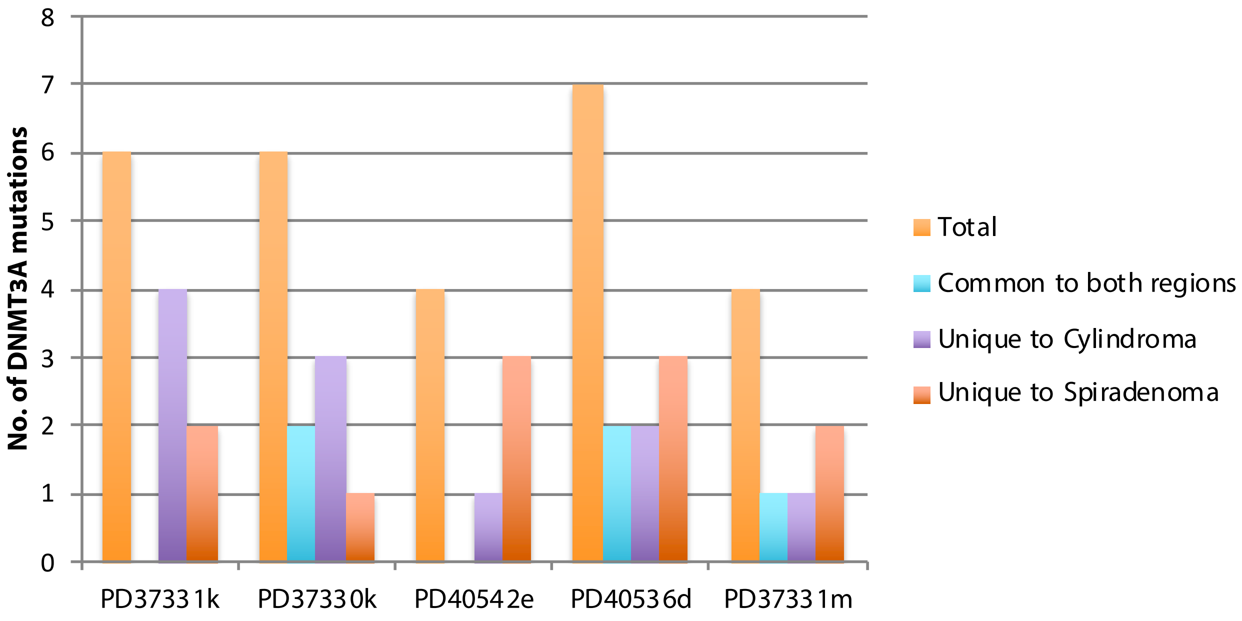


**b**

| Sample | Mutation | Cylindroma VAF | Spiradenoma VAF |
|---|---|---|---|
| PD37331k | p.A70V | 5% | 0% |
| | p.G83R | 5% | 0% |
| | p.I539V | 0% | 2% |
| | p.R635G | 0% | 4% |
| | p.I655T | 7% | 0% |
| | p.I661fs | 11% | 0% |
| PD37330k | p.E61V | 0% | 5% |
| | p.S62G | 3% | 3% |
| | p.S73P | 3% | 1% |
| | p.D92fs | 3% | 0% |
| | p.D668V | 8% | 0% |
| | pR729Q | 43% | 26% |
| PD40542e | p.R23X | 4% | 0% |
| | p.V60E | 0% | 8% |
| | p.V71M | 0% | 5% |
| | p.E205X | 0% | 3% |
| PD40536d | p.K74E | 2% | 6% |
| | c.1123-2A>C | 5% | 21% |
| | p.V622A | 0% | 3% |
| | p.I633fs | 0% | 4% |
| | p.E667V | 8% | 0% |
| | p.G673A | 7% | 0% |
| | p.R729W | 2% | 7% |
| PD37331m | p.K67E | 5% | 0% |
| | p.K299R | 1% | 2% |
| | p.K651R | 0% | 7% |
| | p.C666R | 0% | 7% |

**Supplementary Figure 4. Intratumoral targeted deep sequencing. a** DNA extracted from microdissected regions of 5 tumors with cylindroma and spiradenoma histophenotypes in the same section show the presence of the same driver mutation in 3 out of 5 cases, at varying VAFs, consistent with the contiguous growth of these tumors in three-dimensional space. **b** Table of *DNMT3A* mutations.

**a**

DNMT3A DAPI

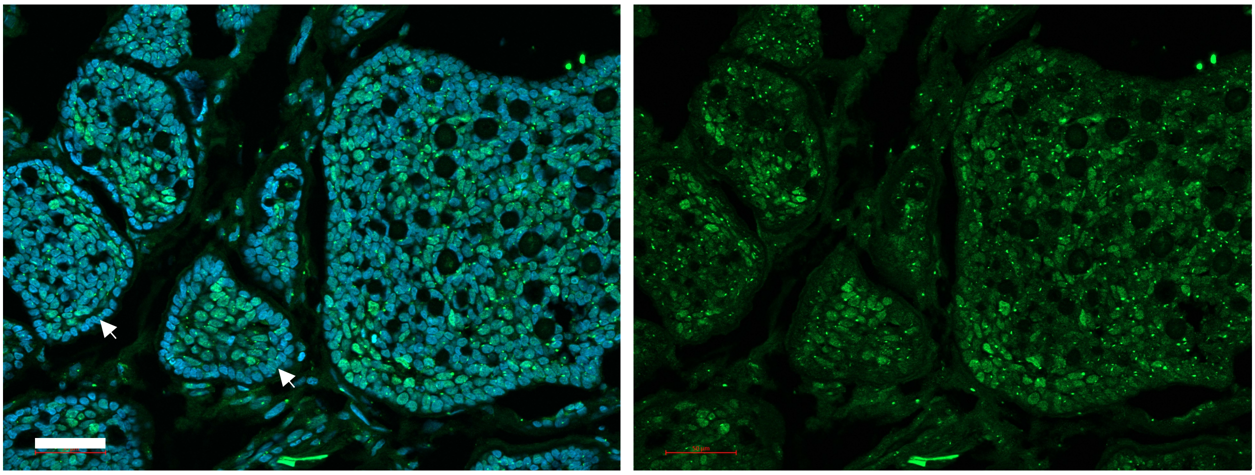

**b** DNMT3A *β*-CATENIN DAPI

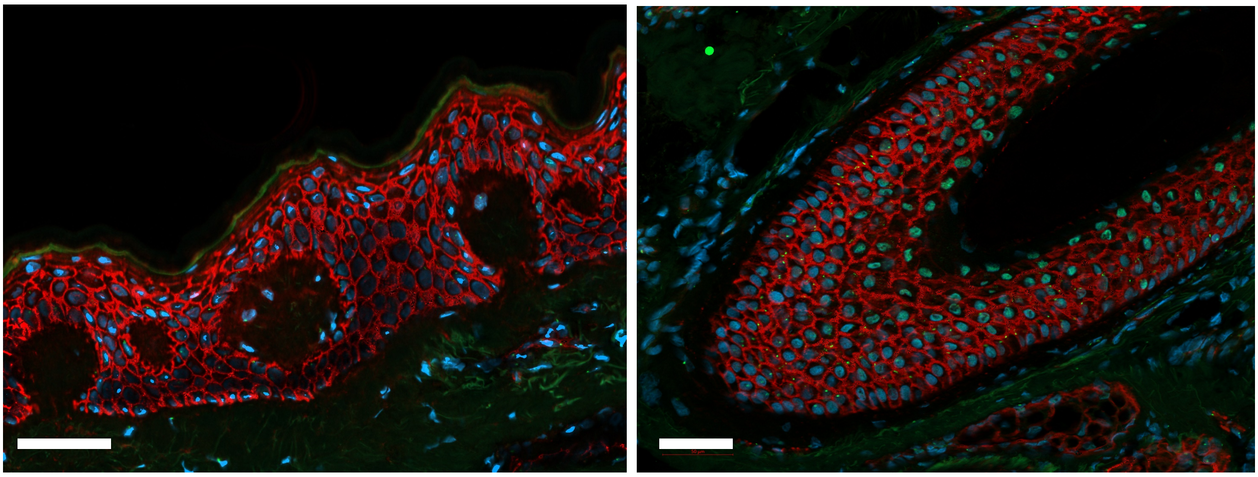

**Supplementary Figure 5. DNMT3A expression in CCS tumors. a** DNMT3A expression is reduced at the periphery in CCS tumor islands (white arrows) compared to central cells. **b** Low DNMT3A expression and membranous beta-catenin expression in control epidermis (left panel) and hair follicles (right panel).

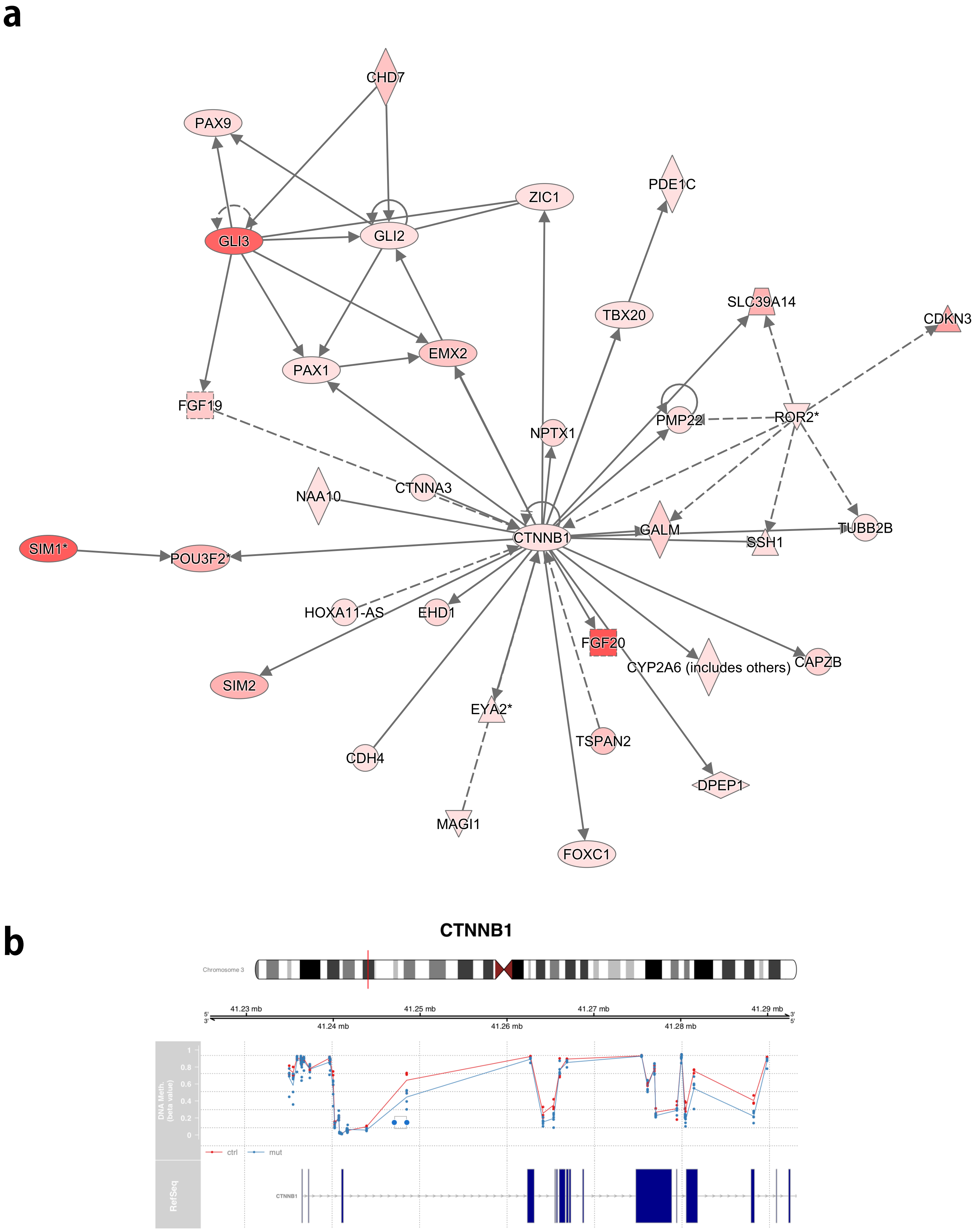


**Supplementary Figure 6. *DNMT3A2* mutated tumors preferentially deregulate methylation of Wnt signalling pathway genes in CCS tumors. a** Pathway analysis of hypomethylated genes in tumors carrying *DNMT3A2* mutant VAF of >0.05 highlighted a network of genes functionally related to ß-catenin, indicated in a network diagram. **b** Methylation profile of beta-catenin is indicated at probe-level.

**a**

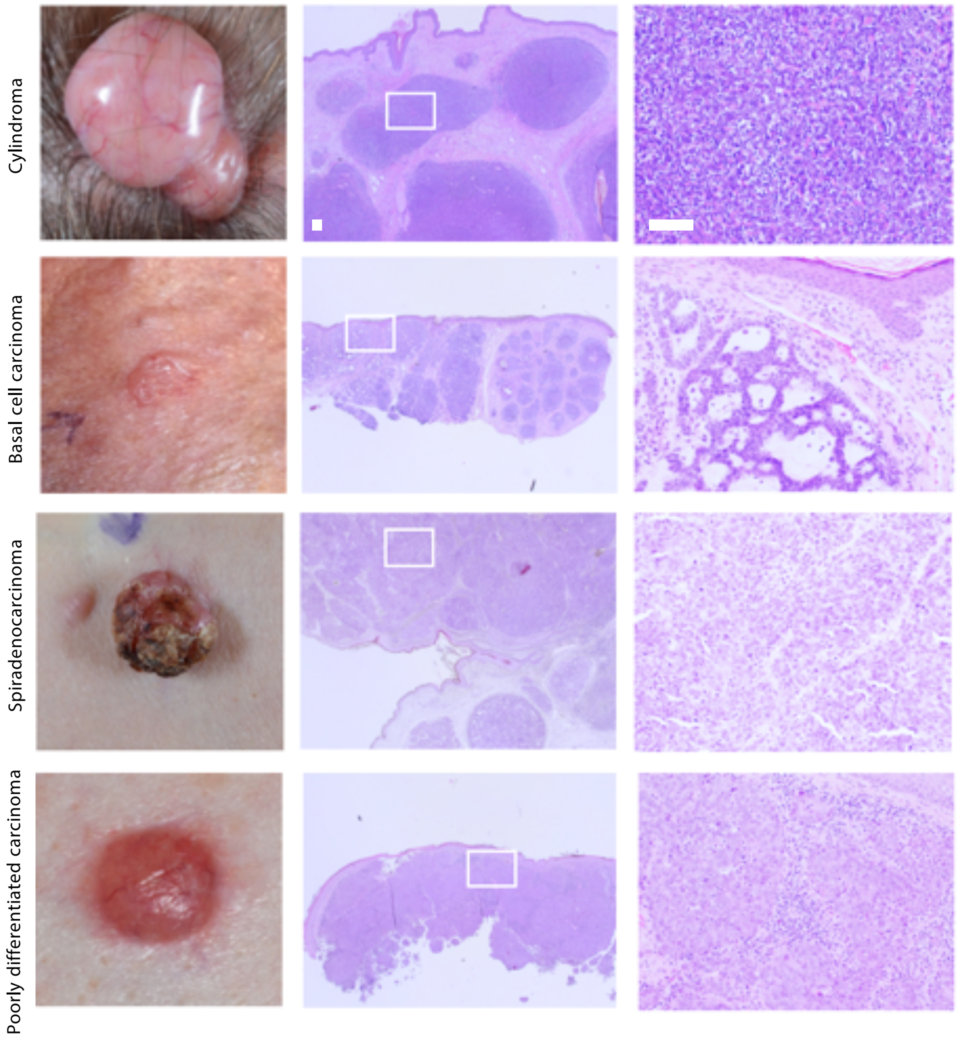


| | *CYLD* | *DRIVERS* |
|---|---|---|
| Cylindroma | *LOH* | *BCOR*<br>*DNMT3A* |
| Basal cell carcinoma | *LOH* | *PTCH1* |
| Spiradenocarcinoma | *LOH* | *MBD4*<br>*CREBBP*<br>*KDM6A*<br>*NOTCH2*<br>*BAP1* |
| Poorly differentiated carcinoma | *No LOH* | *EP300*<br>*TP53* |

**Supplementary Figure 7. Clinical and histological features of malignant tumors seen in CCS. a** Cutaneous tumors seen in CCS, with histological features demonstrated in adjacent panels, with driver mutations summarised. Cylindroma, typically a pink nodule with overlying vessels, demonstrate epithelial cells arranged in cylinders. Malignant spiradenocarcinoma, poorly differentiated adenocarcinoma and basal cell carcinoma demonstrate clinical, histological and genetic differences from this benign tumor.

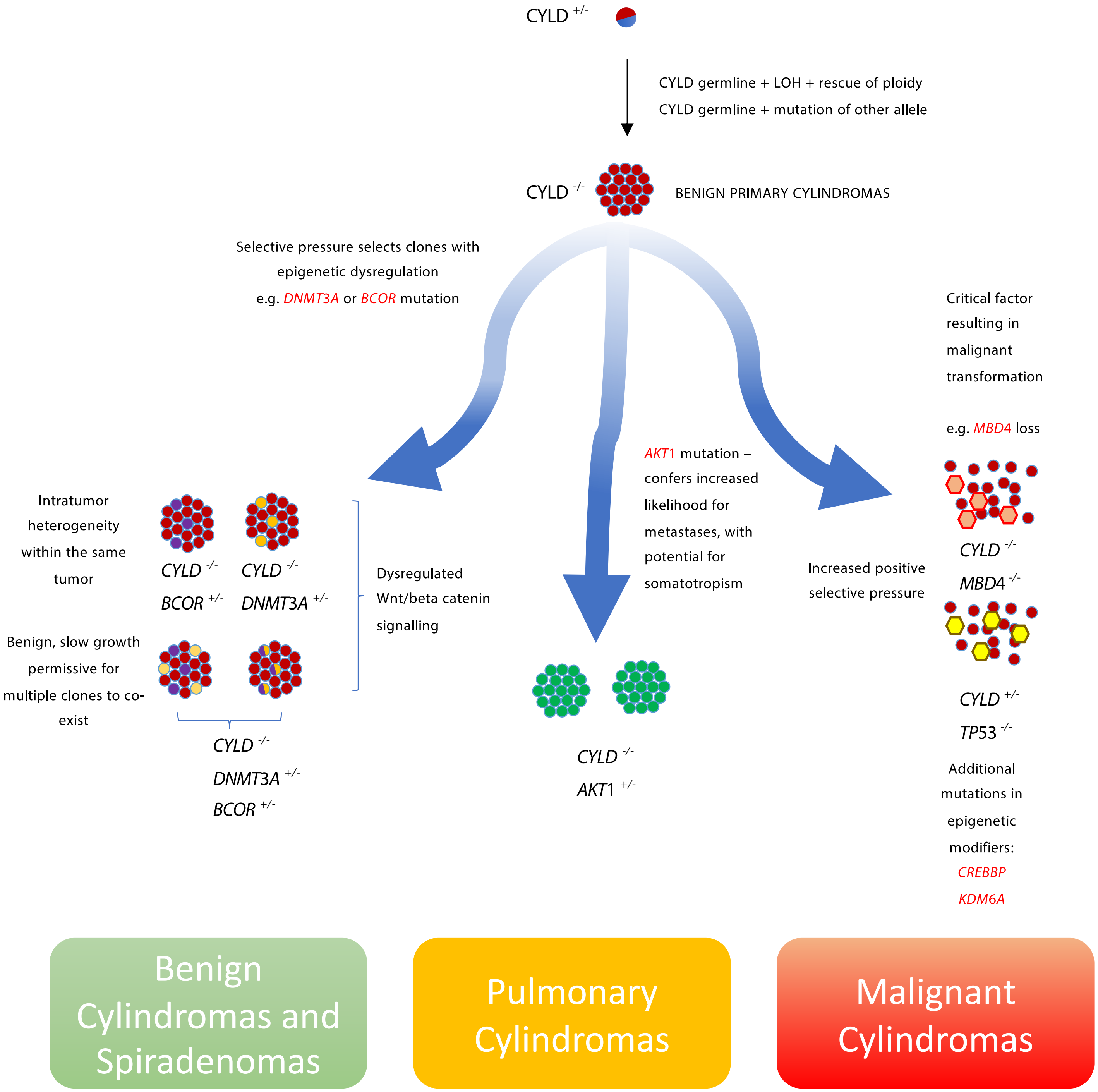


**Supplementary Figure 8. A model of mutation acquisition in CCS tumors.**

| Sample | Patient No | Sex | Age | Genotype | Source | NGS | Array |
|---|---|---|---|---|---|---|---|
| PD36119a | 1 | F | 80 | c.2460delC | Skin | WES | |
| PD36119c | 1 | F | 80 | c.2460delC | Skin | WES, RNAseq | |
| PD37330a | 1 | F | 80 | c.2460delC | Skin | WGS, TDS | EPIC |
| PD37330c | 1 | F | 80 | c.2460delC | Skin | WGS, TDS, RNAseq | EPIC |
| PD37330e | 1 | F | 80 | c.2460delC | Skin | WGS, TDS, RNAseq | EPIC |
| PD37330f | 1 | F | 80 | c.2460delC | Skin | WGS, RNAseq | |
| PD37330g | 1 | F | 80 | c.2460delC | Skin | WGS, TDS | |
| PD37330h | 1 | F | 80 | c.2460delC | Lung | WES | |
| PD37330i | 1 | F | 80 | c.2460delC | Skin | WES, TDS | |
| PD37330j | 1 | F | 80 | c.2460delC | Skin | WES, TDS | |
| PD37330k | 1 | F | 80 | c.2460delC | Skin | TDS | EPIC |
| PD37330l | 1 | F | 80 | c.2460delC | Skin | RNAseq | |
| PD37331a | 2 | F | 64 | c.2460delC | Skin | WGS, RNAseq | |
| PD37331c | 2 | F | 64 | c.2460delC | Skin | WGS | |
| PD37331f | 2 | F | 64 | c.2460delC | Lung | WGS | |
| PD37331g | 2 | F | 64 | c.2460delC | Lung | WGS | |
| PD37331h | 2 | F | 64 | c.2460delC | Lung | WGS | |
| PD37331i | 2 | F | 64 | c.2460delC | Skin | WGS, RNAseq | |
| PD37331j | 2 | F | 64 | c.2460delC | Skin | WES | |
| PD37331k | 2 | F | 64 | c.2460delC | Skin | WES, TDS | |
| PD37331l | 2 | F | 64 | c.2460delC | Skin | WES | |
| PD37331m | 2 | F | 64 | c.2460delC | Skin | TDS | |
| PD37331n | 2 | F | 64 | c.2460delC | Skin | RNAseq | |
| PD37331o | 2 | F | 64 | c.2460delC | Skin | RNAseq | |
| PD40536a | 3 | F | 48 | c.2460delC | Skin | WES | |
| PD40536c | 3 | F | 48 | c.2460delC | Skin | WES | |
| PD40536d | 3 | F | 48 | c.2460delC | Skin | WES, TDS | |
| PD40536e | 3 | F | 48 | c.2460delC | Skin | RNAseq | |
| PD40537a | 4 | M | 84 | c.2469+1G>A | Skin | WES, TDS | |
| PD40537c | 4 | M | 84 | c.2469+1G>A | Skin | WES, TDS | |
| PD40538a | 5 | M | 51 | c.2469 +1G>A | Skin | WES | |
| PD40538c | 5 | M | 51 | c.2469 +1G>A | Skin | WES | |
| PD40538d | 5 | M | 51 | c.2469+1G>A | Skin | RNAseq | |
| PD40538e | 5 | M | 51 | c.2469+1G>A | Skin | RNAseq | |
| PD40538f | 5 | M | 51 | c.2469+1G>A | Skin | RNAseq | |
| PD40539a | 6 | F | 87 | c.2469 +1G>A | Skin | WES | |
| PD40539c | 6 | F | 87 | c.2469 +1G>A | Skin | WES | |
| PD40539d | 6 | F | 87 | c.2469+1G>A | Skin | TDS | EPIC |
| PD40539e | 6 | F | 87 | c.2469+1G>A | Skin | TDS | EPIC |
| PD40540a | 7 | M | 77 | c.2460delC | Skin | WES | |
| PD40540c | 7 | M | 77 | c.2460delC | Skin | WES, TDS | |
| PD40540d | 7 | M | 77 | c.2460delC | Skin | WES | |
| PD40540e | 7 | M | 77 | c.2460delC | Skin | RNAseq | |
| PD40541a | 8 | F | 81 | c.2460delC | Skin | WES, TDS | |
| PD40542a | 9 | F | 75 | c.2460delC | Skin | WES | |
| PD40542c | 9 | F | 75 | c.2460delC | Skin | WES | |
| PD40542d | 9 | F | 75 | c.2460delC | Skin | TDS | EPIC |
| PD40542e | 9 | F | 75 | c.2460delC | Skin | TDS | |
| PD40543a | 10 | F | 43 | c.2460delC | Skin | WES | |
| PD40543c | 10 | F | 43 | c.2460delC | Skin | RNAseq | |
| PD40544c | 11 | F | 57 | c.1112C>A | Skin | WES | |
| PD40544d | 11 | F | 57 | c.1112C>A | Skin | WES | |
| PD40545a | 12 | M | 72 | c.2467C>T | Skin | WES | |
| PD40545c | 12 | M | 72 | c.2467C>T | Skin | WES | |
| PD40546a | 13 | F | 30 | c.1112C>A | Skin | WES | |
| PD40546c | 13 | F | 30 | c.1112C>A | Skin | WES | |
| PD40546d | 13 | F | 30 | c.1112C>A | Skin | WES | |
| PD40547a | 14 | F | 53 | c.2460delC | Skin | RNAseq | |
| PD40548a | 15 | F | 54 | 5.5Mb deletion | Skin | RNAseq | |
| PD40548c | 15 | F | 54 | 5.5Mb deletion | Skin | RNAseq | |
| PD40548d | 15 | F | 54 | 5.5Mb deletion | Skin | RNAseq | |
| PD40548e | 15 | F | 54 | 5.5Mb deletion | Skin | RNAseq | |

**Supplementary Table 1.** Sample list of studied tumours and patient genotypes.

| Sample | Gene | CDS | Protein | Type | Effect | VAF |
|---|---|---|---|---|---|---|
| PD37330h | AKT1 | c.49G>A | p.E17K | Sub | missense | 0.22 |
| PD37331f | AKT1 | c.49G>A | p.E17K | Sub | missense | 0.55 |
| PD37331g | AKT1 | c.49G>A | p.E17K | Sub | missense | 0.44 |
| PD37331h | AKT1 | c.49G>A | p.E17K | Sub | missense | 0.31 |
| PD36119a | BAP1 | c.2036_2055del20 | p.I679fs*31 | Del | frameshift | 0.72 |
| PD37330a | BCOR | c.2606_2607insAGA | p.Y869delins* | Ins | nonsense | 0.31 |
| PD37330h | BCOR | c.1722_1723insC | p.N575fs*36 | Ins | frameshift | 0.087 |
| PD37330i | BCOR | c.1061_1062insAA | p.Y354fs*1 | Ins | frameshift | 0.378 |
| PD37330j | BCOR | c.2221_2225delGAGAAinsTTTC | p.E741fs*45 | Complex | frameshift | 0.309 |
| PD40537a | BCOR | c.1059delC | p.Y354fs*24 | Del | frameshift | 0.074 |
| PD40540c | BCOR | c.3621_3622insA | p.Q1208fs*8 | Ins | frameshift | 0.075 |
| PD40543a | BCOR | c.2428C>T | p.R810* | Sub | nonsense | 0.048 |
| PD40545a | BCOR | c.1572_1573insTGGCAAAAGC | p.M525fs*35 | Ins | frameshift | 0.350 |
| PD36119a | CREBBP | c.4337G>A | p.R1446H | Sub | missense | 0.31 |
| PD36119a | CREBBP | c.3307C>T | p.R1103* | Sub | nonsense | 0.39 |
| PD37330a | DNMT3A | c.1518C>G | p.H506Q | Sub | missense | 0.36 |
| PD37330a | DNMT3A | c.1517A>G | p.H506R | Sub | missense | 0.37 |
| PD37330c | DNMT3A | c.1531G>A | p.G511R | Sub | missense | 0.42 |
| PD37330g | DNMT3A | c.2339T>C | p.I780T | Sub | missense | 0.34 |
| PD40536d | DNMT3A | c.1123-2A>C | p.? | Sub | ess_splice | 0.095 |
| PD40537a | DNMT3A | c.2644C>T | p.R882C | Sub | missense | 0.26 |
| PD40541a | DNMT3A | c.2638delA | p.M880fs*1 | Del | frameshift | 0.182 |
| PD40536c | EP300 | c.584C>A | p.S195* | Sub | nonsense | 0.085 |
| PD40536c | EP300 | c.2855C>G | p.S952* | Sub | nonsense | 0.082 |
| PD36119a | KDM6A | c.1972C>T | p.R658* | Sub | nonsense | 0.65 |
| PD36119a | NOTCH2 | c.6909_6910insC | p.I2304fs*9 | Ins | frameshift | 0.37 |
| PD40544c | PTCH1 | c.843_863del21insG | p.L282fs*30 | Complex | frameshift | 0.320 |
| PD40536c | TP53 | c.614_615insGTA | p.E204_Y205ins* | Ins | nonsense | 0.312 |

**Supplementary Table 2.** Variant allele fraction of coding mutations detected using whole genome sequencing and whole exome sequencing.

| Sample | Source | Mutant reads | Total reads | Variant Allele Fraction |
|---|---|---|---|---|
| PD37330a | Cylindroma | 21 | 45 | 0.47 |
| PD37330c | Cylindroma | 15 | 37 | 0.41 |
| PD37330d | Normal skin | 12 | 31 | 0.39 |
| PD37330e | Cylindroma | 20 | 35 | 0.57 |
| PD37330f | Cylindroma | 22 | 50 | 0.44 |
| PD37330g | Cylindroma | 25 | 45 | 0.56 |
| PD37331a | Cylindroma | 19 | 44 | 0.43 |
| PD37331c | Cylindroma | 19 | 38 | 0.50 |
| PD37331d | Normal skin | 18 | 43 | 0.42 |
| PD37331e | Normal lung | 19 | 37 | 0.51 |
| PD37331f | Pulmonary cylindroma | 24 | 43 | 0.56 |
| PD37331g | Pulmonary cylindroma | 22 | 40 | 0.55 |
| PD37331h | Pulmonary cylindroma | 18 | 36 | 0.50 |
| PD37331i | Cylindroma | 20 | 46 | 0.43 |
| Exome samples | | | | |
| PD36119a | Malignant cylindroma | 166 | 202 | 0.82 |
| PD36119c | Cylindroma | 81 | 153 | 0.53 |

**Supplementary Table 3.** *MBD4* somatic status in pedigree 1 tumors.

**Supplementary Table 4.** A full list of all mutations detected is available at https://www.dropbox.com/sh/iyuiyk666fc4v1g/AAAxQni30RARao4RPqts42Kaa?dl=0
